# Multiphase condensates from a kinetically arrested phase transition

**DOI:** 10.1101/2022.02.09.479538

**Authors:** Nadia A. Erkamp, Tomas Sneideris, Hannes Ausserwöger, Daoyuan Qian, Seema Qamar, Jonathon Nixon-Abell, Peter St George-Hyslop, Jeremy D. Schmit, David A. Weitz, Tuomas P.J. Knowles

## Abstract

The formation of biomolecular condensates through liquid-liquid phase separation from proteins and nucleic acids is emerging as a spatial organisational principle used by living cells. Many such biomolecular condensates are not, however, homogeneous fluids, but contain an internal structure consisting of distinct sub-compartments with different compositions. In many instances, such compartments inside the condensate are depleted in the biopolymers that make up the condensate. Here, we describe that this multiphase structure arises from a kinetically arrested phase transition. The combination of a change in composition coupled with a slow response to this change can lead to the spontaneous formation of multiple emulsions consisting of several inner cores within a polymer-rich middle phase. In the case of liquid-like biomolecular condensates, the slow diffusion of biopolymers causes nucleation of biopolymer-poor liquid inside of the condensate to achieve the new equilibrium composition. This framework shows that multiphase condensates can be a result of kinetic trapping, rather than thermodynamic stability, and provides and avenue to understand and control the internal structure of condensates *in vitro* and *in vivo*.

## Introduction

It is crucial for life to have spatial and temporal control over the biochemical reactions it performs.^1,2^ Membraneless organelles in the form of biomolecular condensates are an important tool to organize proteins and nucleic acids and they perform a range of different functions, including storage^3^, processing^4^, signalling^5^ and modulating gene expression^6^. Crucially, many functional condensates, such as stress granules^7^, paraspeckles^8^, nuclear speckles^9^ and the nucleolus^10^, have an internal structure; they consist of sub-compartments containing distinct compositions of proteins and nucleic acids. Interestingly, such sub-compartments that are poor in the biopolymers that compose the condensate have been widely observed.^11–15^. A similar “core-shell” structure has also been observed in microgels^16^. Moreover, in living cells, TDP-43 rich droplets with nucleoplasm-filled vacuoles,^17^ nuclear and cytoplasmic germ granules with hollow centers,^18–20^ and poly(A) RNA nuclear condensates with low-density space have been observed.^21^ While this class of structure is thus commonly encountered in nature, it is not yet clear what the driving forces are for the formation of biopolymer-poor phases inside of the biopolymer-rich condensate. Here, we show that this multiphase structure forms due to a kinetically arrested phase transition when the condensate is out-of-equilibrium. Limited diffusion causes condensates to deviate from the binodal during composition changes, causing nucleation of dilute phase inside of the condensates. The deviation from equilibrium can be induced by changes in the environment and solidification of the condensates.

## Results and discussion

### Cavity formation is reversible with composition change

We explore the mechanism behind this multiphase structure with condensates formed from PolyA RNA and PEG. These condensates don’t significantly solidify over time and change composition with temperature, containing more PolyA at 20 °C than at 55 °C and dissolving at approximately 65 °C (Supplementary Fig. 1). We vary the temperature to determine the effect of changing the condensate composition on the formation of cavities, liquids poor in PolyA and PEG inside the condensate (Supplementary Fig. 2). We observe these cavities in condensates at 20 °C, but not at 55 °C (Fig. 1A). This is the case for multiple cycles of heating and cooling to these temperatures, implying the cavities are created and removed in between these temperatures (Supplementary Fig. 3 and 4). We check this, starting with a condensate with cavities at 20 °C (Fig. 1B.1). When heating to 30 and 55 °C, we observe that cavities become smaller and completely disappear respectively (Fig. 1B.2-5 and Supplementary Fig. 5). After the cavities are completely removed, we lower the temperature of these condensates and cavities form again (Fig. 1B.6). The cavities grow with decreasing temperature (Fig. 1B.7-9). At 20 °C, we obtain condensates that look very similar to the ones before the temperature change (Fig. 1B.10). We conclude that cavity formation is reversible and takes place during a composition change. This could suggest that cavity formation is a kinetic process, in which case cavity formation would depend strongly on the rate at which the composition changes.

**Fig. 1:**
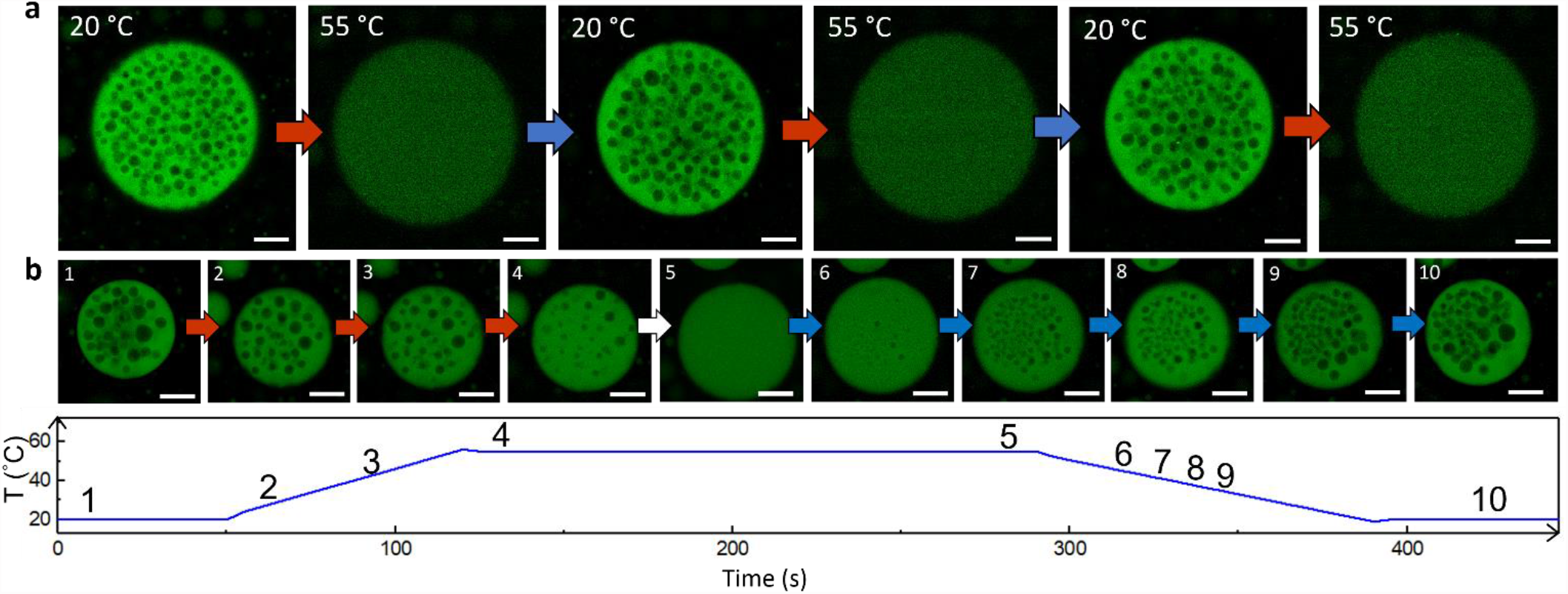
Temperature-dependent and reversible cavity formation. **a** Confocal microscopy images of multiple cycles of cavity formation and removal (Supplementary Fig. 3). Intensity differences are to show that the concentration of PolyA in the dense phase is higher at 20 than at 55 °C (Supplementary Fig. 4). **b** Confocal microscopy images of temperature-induced morphology changes of PolyA-PEG condensates. The intensity of the PolyA-rich phase is kept constant at every temperature and is thus not an indication of PolyA concentration. All scale bars represent 25 μm.

### Cavity formation is a kinetic process

To probe whether cavities originate from a thermodynamic or kinetic process, we varied the rate at which condensates are cooled from 55 to 20 °C. A condensate with a radius of 24 μm will contain only 1 cavity after cooling slowly at 1 °C/min, but 24 cavities after cooling faster at 20 °C/min (Fig. 2, Supplementary Fig. 6A). Thus, depending on the rate of composition change, we obtain a kinetic product, a condensate with many cavities, or a more thermodynamically stable product, a condensate with less cavities. This observation implies that cavities are formed when the condensate is unable to reach the thermodynamic equilibrium. If this is the case, we would expect that the number of cavities also depends on the size of the condensate, as smaller condensates might equilibrate faster with the surrounding dilute phase than large condensates. Indeed, a condensate with a radius of 14 μm does not form cavities at any of the tested cooling rates and the condensate with a radius of 38 μm contains more cavities than the condensate with a 24 μm radius, confirming cavity formation is a kinetic process (Supplementary Fig. 6B). Condensates with cavities might be higher in energy due to the additional surface area from the cavities, which increases the surface energy. If this kinetic product was indeed higher in energy, we could expect that over time the number of cavities in condensates will be reduced.

**Fig. 2:**
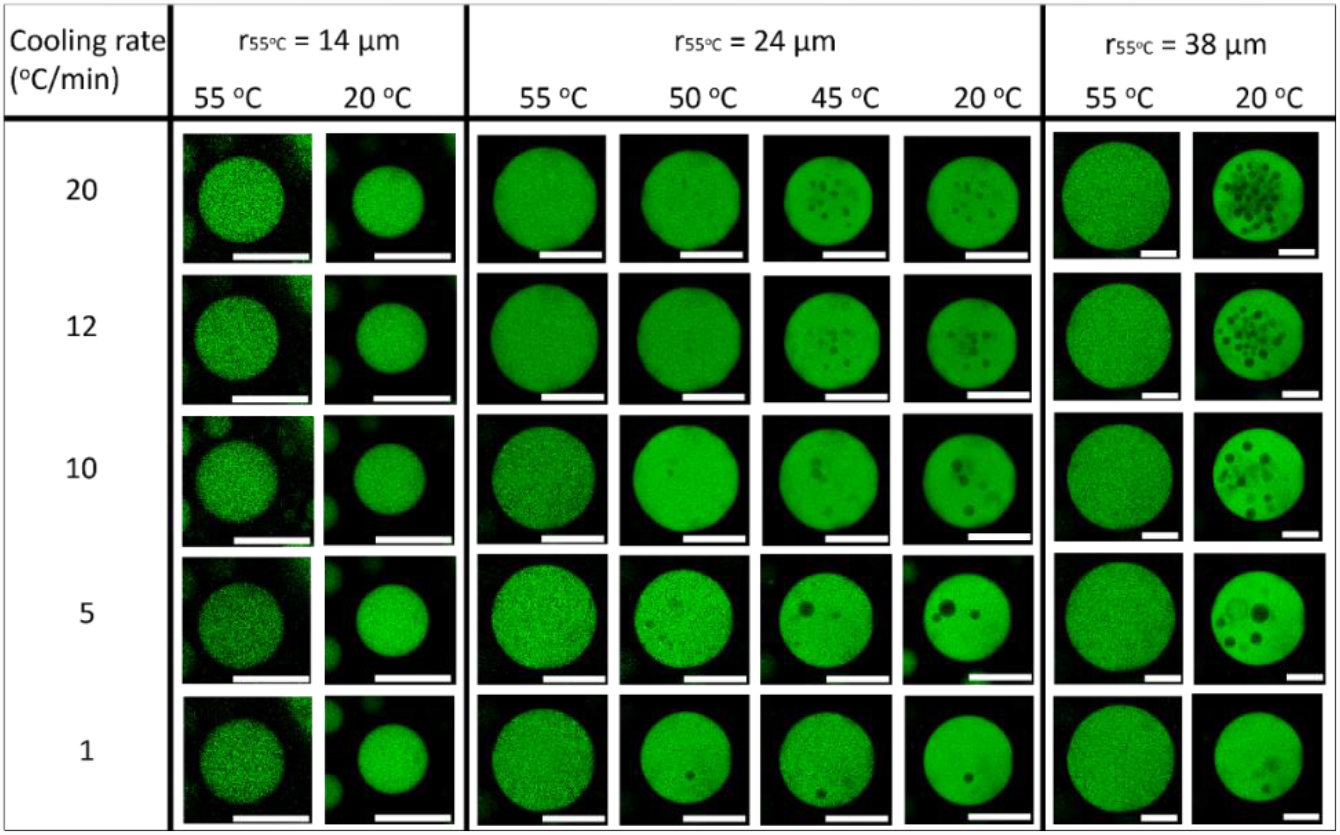
Formation of cavities depends on cooling rate or composition change rate, and condensate size. Confocal microscopy images of condensates with a radius of 14, 24 and 38 μm being cooled from 55 to 20 °C at different rates. The number of cavities formed during cooling depends both on the size of the condensate and on the rate at which the condensates is cooled (Supplementary 6). All scale bars represent 25 μm.

### Cavities are similar in composition to the bulk dilute phase

We observe a condensate with cavities over time (Fig. 3A). Cavities move through the condensate and merge with each other once coming into contact. Similarly, cavities merge with the dilute phase surrounding the condensate, indicating that the liquid in the cavities and surrounding dilute phase are similar. Over time, the number of cavities decreases via these two mechanisms, as they form less, but larger cavities or “escape” to the outer dilute liquid, decreasing surface energy (Supplementary Fig. 6C). We study these merging events and the merging of the condensates to confirm that the cavities are similar in composition to the surrounding dilute phase. The time it takes for two droplets in contact to merge into one larger spherical droplet depends on the capillary velocity, which is a function of the surface tension and the viscosity.^6,22,23^ To probe whether the cavity/condensate phase surface tension differs from that of the dilute phase/condensate face interface, we evaluated the capillary velocities corresponding to the three classes of fusion events. Since the viscosity entering the capillary velocity equation is always that of the dense condensate phase, any differences show differences in the surface tensions between the different interfaces. To evaluate the capillary velocity, we measured the aspect ratio, the length and width of the two merging droplets as a function of time. The plot of aspect ratio against time is then fitted with the exponential decay function to find the characteristic merging time τ.^6,23^ A plot of this merging time as a function of the size of the merging droplets gives the inverse capillary velocity.^6,22,23^ For merging condensates, merging cavities and escaping cavities, we find very similar values for inverse capillary velocities (Fig. 3B, C, D). Thus, the surface tension between the condensate and the outer dilute phase, and the condensate and the cavity is very similar, implying that cavities are dilute phase liquid trapped in the condensate.

**Fig. 3:**
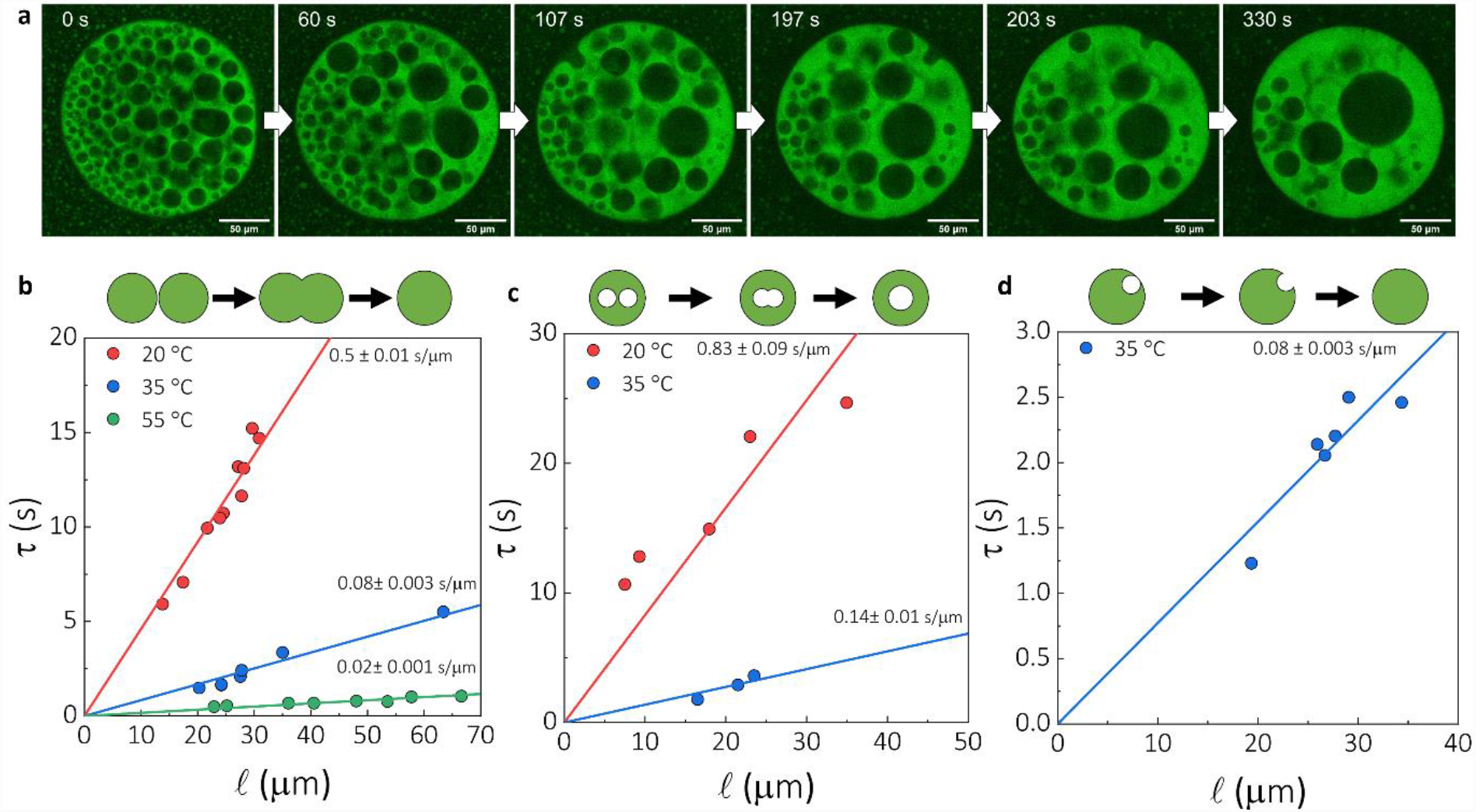
Dynamics of cavities and merging events. **a** Cavities inside of PolyA-PEG condensates can merge with each other and with the outer dilute phase. The sample was incubated at 35 °C. From the merging of **b** condensates, **c** cavities and **d** cavities with the outer dilute phase, we extract the characteristic merging time τ. The slope of the plot τ against the average diameter of merging droplets, ℓ, is the inverse capillary velocity, which is approximately equal to the viscosity over the surface tension. ^6,22^

### Cavities form due to a kinetically arrested phase transition

The binodal curve of a phase diagram describes the equilibrium concentrations in the dilute and dense phase for a phase-separated system. When a change in the environmental conditions occurs, the composition of the dilute and dense phase will change to their new equilibrium positions on the binodal curve. We have observed in our PolyA-PEG model system that if a composition change occurs relatively quickly, the system can’t equilibrate and cavities are formed. We conclude that during the composition change, the system deviates from the binodal curve and droplets of the dilute phase can nucleate inside of the condensate (Fig. 4A, Supplementary Fig. 7 and Supplementary text 1 and 2). Notably, the spinodal and binodal are in the opposite position from what is often shown in phase diagrams, since this figure shows the emergence of the dilute phase inside of a dense phase. Cavities form because the condensate is unable to stay on the binodal curve, causing nucleation of dilute phase. Condensates typically have a low surface tensions,^24^ which leads to a low energy penalty for cavity formation and thus a low nucleation barrier for forming a multiphase condensate. The slow response to composition changes could be caused by the viscoelasticity of the condensate, as it is unable to change to the size required for the composition change to complete, or could be due to slow diffusion rates, preventing the condensate from reaching the ideal concentrations during the composition change.

**Fig. 4:**
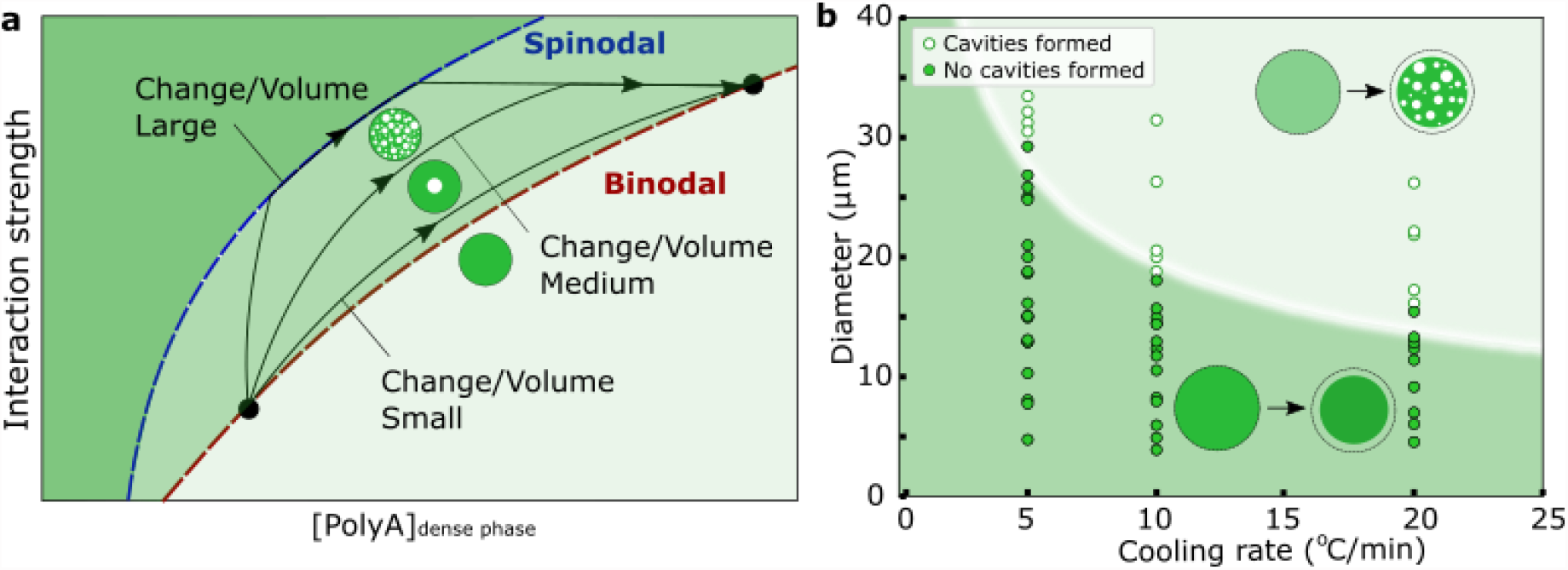
Mechanism of cavity formation. **a** The ideal biopolymer concentration in the condensate is increased. Depending on factors like the rate of the change and the size of the condensate, the condensate can follow the binodal curve quite closely or deviate from the binodal curve, at which point biopolymer-poor droplets can nucleate inside of the condensates. **b** Experimental data on the critical condensate size for the nucleation of cavities (points) are fit with a scaling law to give an effective diffusion coefficient of 12*μm*^2^/*s* shown as the curve in the background. The ability to from cavities depends strongly on the size of the condensate and the rate of composition change.

### Slow diffusion can cause cavity formation

To determine whether viscoelasticity could be a limiting factor, we measured the volume of a condensate when cooling at 1 and 20 °C/min. We found no significant difference between the volumes indicating that at both cooling rates the equilibrium volume can be reached and viscoelasticity is unlikely the cause of the slow response (Supplementary Fig. 8). This matches with the liquid-like, rather than gel-like, nature of these condensates. To determine if diffusion of biopolymers in the condensate could be a limiting factor for equilibration during quick composition changes, we determined the diffusion coefficients of PolyA and PEG in the condensates. Using fluorescence recovery after photobleaching (FRAP)^25^, we found that diffusion through these dense condensates is indeed slow (Supplementary Fig. 9). Additionally, we determined the effective diffusion coefficient *D*_*eff*_ by measuring the critical size of condensates for cavity formation (Fig. 4B). We fit this data using the scaling law 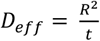 derived from the diffusion equation,^26,27^ in which R is the diffusion length scale or radius of the condensate and 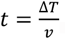, where *v* is the cooling rate and Δ*T* ≈ 5 °C is the amount of temperature change needed to form cavities. Fitting this scaling law gives us an effective diffusion coefficient of approximately 12*μm*^2^/*s*, which is a very similar value to the diffusion coefficients measured using FRAP. We conclude that the slow diffusion of macromolecules in liquid-like biomolecular condensates is the limiting factor in the condensates response to environmental changes. This raises the question if cavity formation is specific to liquid-like condensates, or if this could also occur in more gel-like materials.

### Kinetically arrested phase transitions in diverse systems

Reconstituted stress granules, which are similar in composition to the condensates found in cells,^28^ can form cavities as a consequence of a composition change (Fig. 5A). Interestingly, as they age and solidify, cavities also form and merge, despite environmental conditions staying the same (Fig. 5B). From this we conclude that cavities can be produced in more gel-like materials and that the solidification causes the composition change. The slow response could be caused by a combination of limited diffusion and viscoelasticity. Notably, cavities are unable to merge with the surrounding dilute phase, resulting in condensates that look “hollow”, also in the case of antibody-DNA condensates (Fig. 5C). This might be caused by a gelled outer shell on the condensate, preventing dilute phase droplets from passing through. In gels, cavities can be formed during syneresis, in which a gel changes composition^29–31^, and during a change in temperature.^16^ Thus, cavities can be formed in liquid- and gel-like materials as a consequence of a composition change. The cause of this composition change can be temperature change, addition of RNA or solidification of the material. In literature, cavities have been observed as a consequence of these factors,^11–13^ but also as a consequence of pH change or addition of salt, also factors influencing the composition of condensates.^14,15^ Our kinetically arrested phase transition framework allows us to understand how a multiphase structure can arise in many different systems due to kinetic trapping, rather than thermodynamic stability. This work highlights the importance of considering kinetics to understand the function and structure of condensates, particularly in cells which operate out-of-equilibrium.

**Fig. 5:**
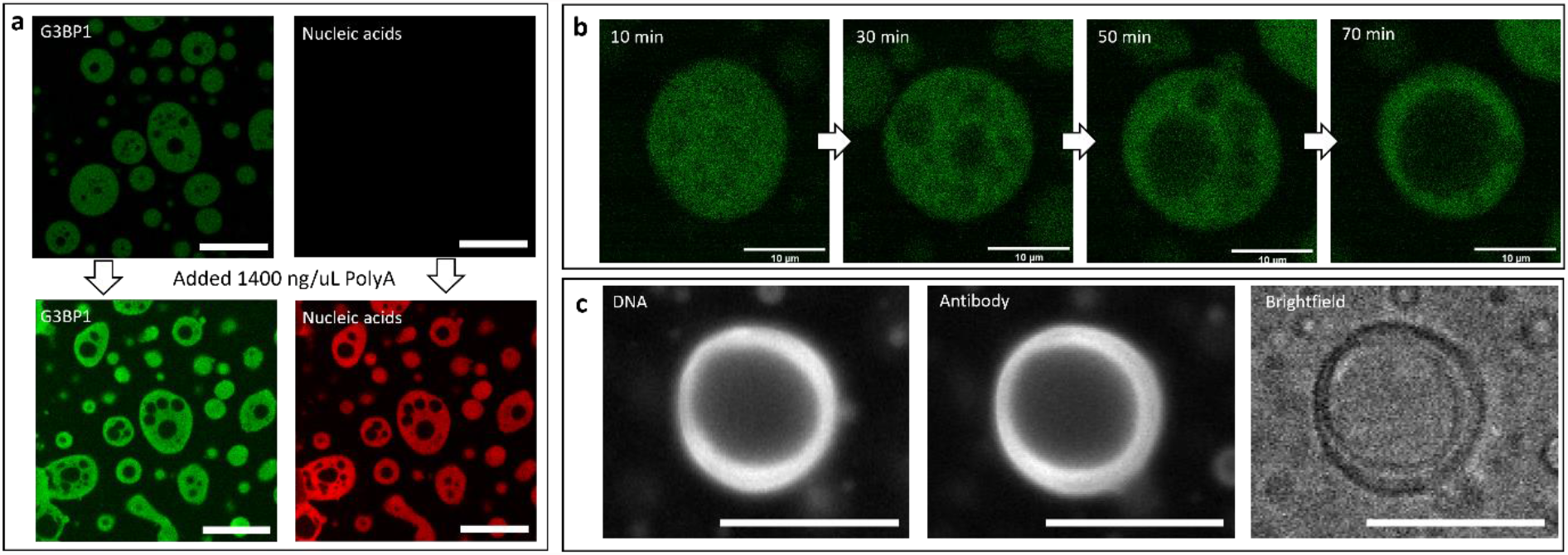
Cavity formation in more gel-like condensates. **a** Reconstituted stress granules can from cavities due to composition change, induced by adding PolyA RNA to the lysate solution. **b** Additionally, cavity formation can also be induced by solidification of the condensates. **c** Cavity formation over time was also observed in HzATNP v.4 Antibody-DNA condensates, with the widefield pictures shown taken after 1 hour at room temperature. Cavities in solidified condensate are unable to merge with the surrounding dilute phase. Unless stated otherwise, all scale bars represent 25 μm.

## Conclusion

Biomolecular condensates, including those found in cells, often contain distinct sub-compartments with different compositions. Sub-compartments poor in biopolymers are common, but the mechanism underlying their formation was not yet known. We have presented that these subcompartments are formed due to a composition transition with a slow response leading to a kinetically arrested phase transition. In condensates, slow diffusion of biopolymers causes the condensate to deviate from the binodal curve, causing the nucleation of biopolymer-poor droplets in the condensate. This mechanism allows us to understand and control the formation biopolymer-poor liquids inside of condensates using a kinetic framework.

## Supporting information

Supplementary figures 1-9, supplementary texts 1 and 2 and method section

## Acknowledgements

We thank Prof. Frans Spaepen for discussions. The research leading to these results has received funding from the Royall Scholarship (N.A.E.), the European Union’s Horizon 2020 research and innovation programme under the Marie Skłodowska-Curie grant MicroREvolution (agreement no. 101023060; T.S.), Global Research Technologies Novo Nordisk A/S (H.A, T.P.J.K.), a Henry Wellcome Fellowship (218651/Z/19/Z, J.N.A.), Canadian Institutes of Health Research (Foundation Grant and Canadian Consortium on Neurodegeneration in Aging Grant, P. St G.H.), Wellcome Trust Collaborative Award 203249/Z/16/Z (P. St G.H., T.P.J.K.), and US Alzheimer Society Zenith Grant ZEN-18-529769 (P. St G.H.), the NIGMS (R01GM141235; J.D.S.), the European Research Council under the European Union’s Seventh Framework Programme (FP7/2007-2013) through the ERC grants PhysProt (agreement no. 337969; T.P.J.K.) and the Newman Foundation (T.P.J.K., T.S.).

## Author Contributions

N.A.E., T.S., J.S., H.A. and T.P.J.K. conceived the study. N.A.E., T.S., D.A.W., H.A., D.Q. and J.N.-A. performed investigation. S.A., T.P.J.K. and P.St.G-H. provided resources. T.P.J.K. and P.St.G-H. acquired funding. N.A.E., T.S. and T.P.J.K. wrote the original draft, all authors reviewed and edited the paper.

