## Supplementary figures 1-9, supplementary texts 1 and 2 and method section for "Multiphase condensates from a kinetically arrested phase transition"

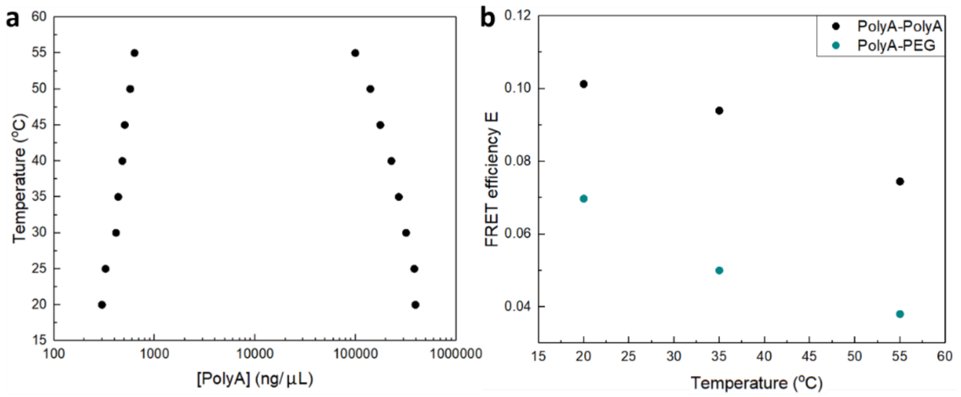


**Supplementary Fig. 1: The composition of PolyA-PEG condensates changes with temperature. a** Approximate PolyA concentration in the dilute and dense phase determined using confocal microscopy as a function of temperature. Notably, while measurements are corrected for the effect that temperature has on dye brightness (Supplementary Fig. 4), they are still approximations, since changes in the quantum efficiency of the dye due to differences in environments are difficult to correct for. The PolyA concentration in the dilute phase match with the concentrations we determined by spinning down the condensates and measuring the concentration in the dilute phase using a nanodrop machine. **b** FRET-FLIM (Forster Resonance Energy Transfer - Fluorescence-lifetime imaging microscopy) experiments show decreasing distance between the biopolymers in PolyA-PEG condensates with temperature. The FRET efficiency, E, is determined using $E = 1 - \frac{\tau_{DA}}{\tau_D}$, where $\tau_{DA}$ is the lifetime of the donor dye in the presence of the acceptor dye and $\tau_D$ is the lifetime of the donor dye without the acceptor. The FRET efficiency is inversely correlated to the distance between the molecules, confirming that temperature causes a composition change in PolyA-PEG condensates.

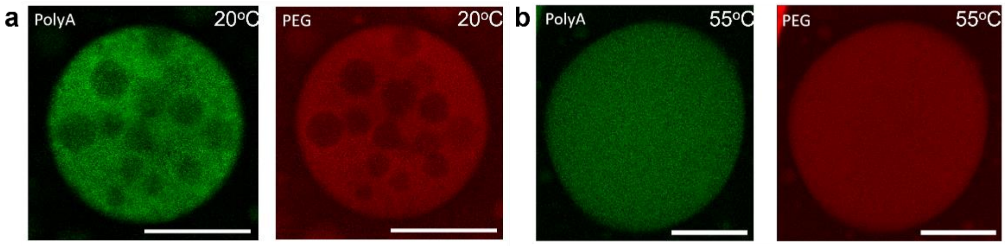


**Supplementary Fig. 2: PolyA-PEG condensate at 20 and 55 °C. a** A condensate with cavities is formed by cooling it at 20 °C/min from 55 to 20°C. Both the PolyA and PEG are labelled showing that the cavities are poor in both PolyA and PEG, like the surrounding dilute liquid. **b** When heating this condensate to 55 °C, the condensate expands. The condensate is still enriched in PolyA and PEG in comparison to the bulk dilute liquid. Scale bars represent 25 µm.

*[1]Yusuf Hamied Department of Chemistry, Centre for Misfolding Diseases, University of Cambridge, Lensfield Road, Cambridge, CB2 1EW, UK. [2]Cambridge Institute for Medical Research, Department of Clinical Neurosciences, University of Cambridge, Cambridge CB2 0XY, UK. [3]Department of Medicine (Division of Neurology), University of Toronto and University Health Network, Toronto, Ontario M5S 3H2, Canada. [4]Department of Neurology, Columbia University, 630 West 168th St , New York, NY 10032, USA. [5]Department of Physics, Kansas State University, Manhattan, KS 66506, USA. [6]Department of Physics, Harvard University, 17 Oxford Street, Cambridge, MA 02138, USA. [7]John A. Paulson School of Engineering and Applied Sciences, Harvard University, Cambridge, MA 02138, USA. [8]Wyss Institute for Biologically Inspired Engineering, Harvard University, Cambridge, Massachusetts 02138, USA. [9]Cavendish Laboratory, Department of Physics, University of Cambridge, J J Thomson Ave, Cambridge, CB3 0HE, UK. †These authors contributed equally. *Email:*

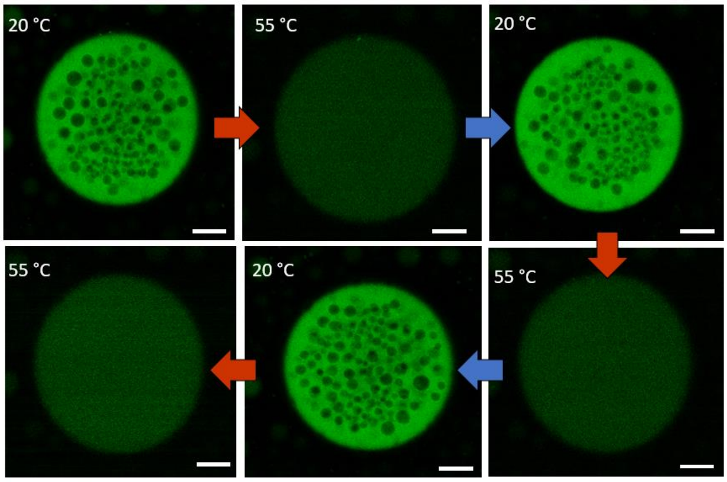


**Supplementary Fig. 3: Temperature dependent reversible cavity formation in PolyA-PEG condensates.** After cavities have been formed for 3 cycles (Fig. 1a), the formation and removal of BPPs can take place for at least another 3 cycles). The dye intensity has been corrected for intensity changes as a result of temperature differences (Supplementary Fig. 4). All scale bars represent 25 µm.

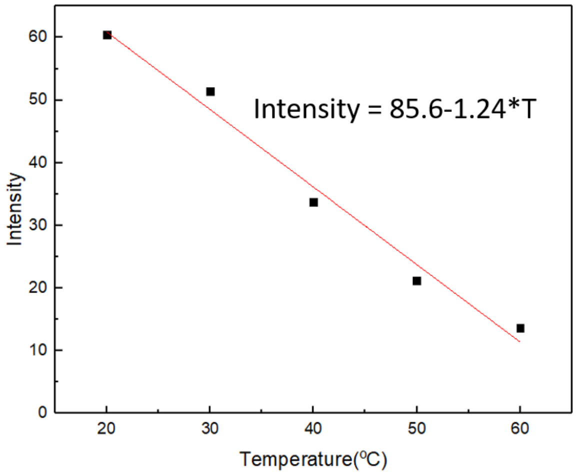


**Supplementary Fig. 4: Correcting the dye intensity of BactoView™ at different temperatures.** The intensity of a solution containing PolyA and BactoView™ (25×) was measured at 20, 30, 40, 50 and 60 °C. From fitting the intensity as a function of temperature, we find that the BactoView™ dye is at 55 °C 29% as bright as it is at 20 °C. To correct for this effect, pixel values in pictures at 55 °C have been increased by ($\frac{100}{29}$ =) 3.45.

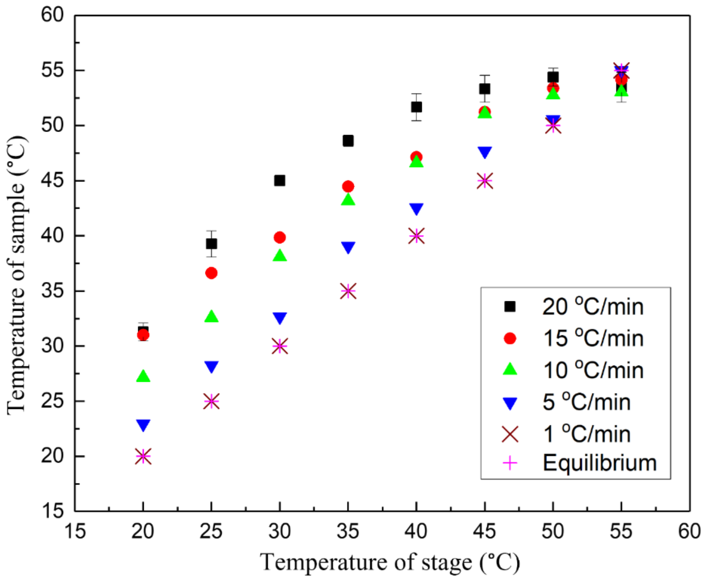


**Supplementary Fig. 5: Conversion of the temperature readout of the temperature control stage to the temperature in the sample.** The samples in figure 1 and 2 of the main text were cooled from 55 to 20 °C at different rates. To achieve this, the samples are placed on a stage, which displays the temperature of the stage itself. We have used the temperature-dependent brightness of AF647 to determine the temperature of the sample in comparison to the one displayed on the stage to correctly report the temperatures and rates shown in figure 1 and 2. For rates faster than 1 °C/min, the sample temperature will lag behind that of the stage, but the same rate is achieved, expect for the rate of 15 °C/min, in which case the temperature change in the sample was 12 °C/min.

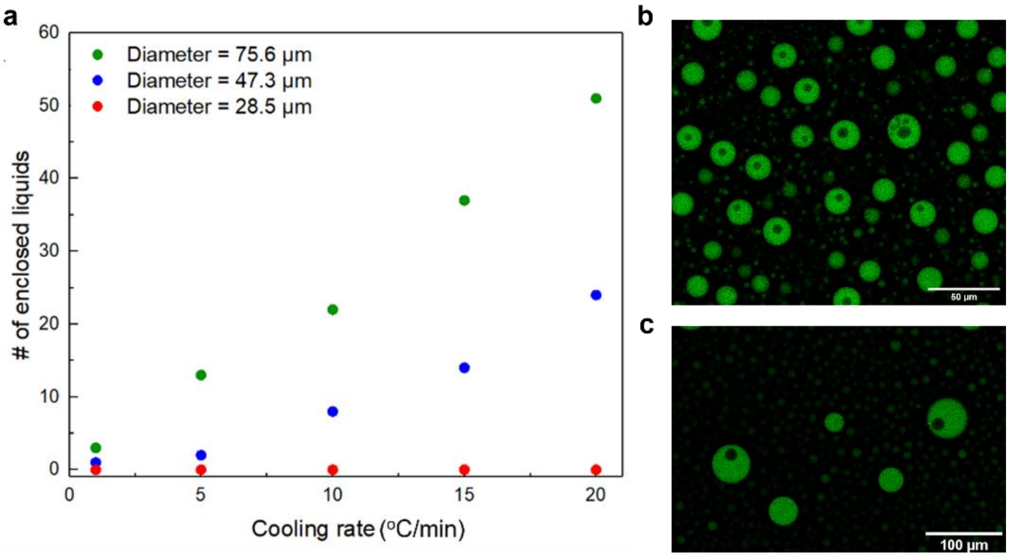


**Supplementary Fig. 6: Number of cavities depends on condensate size, cooling rate and time since formation. a** Different sizes PolyA-PEG condensates are cooled at different rates from 55 to 20°C showing more cavities are formed in larger condensates, which are cooled quicker. Notably, these numbers were determined by analysing a z-stack of pictures of the entire condensate, not just the slice shown in figure 2. **b** Confocal microscopy picture of condensates cooled at 20 °C/min showing that cavity formation requires condensates of sufficient size. Condensates significantly larger than this size can form multiple cavities. **c** Condensates after 100 hours at 20 °C contain much less cavities than right after formation, since cavities can merge with each other and the surrounding dilute phase.

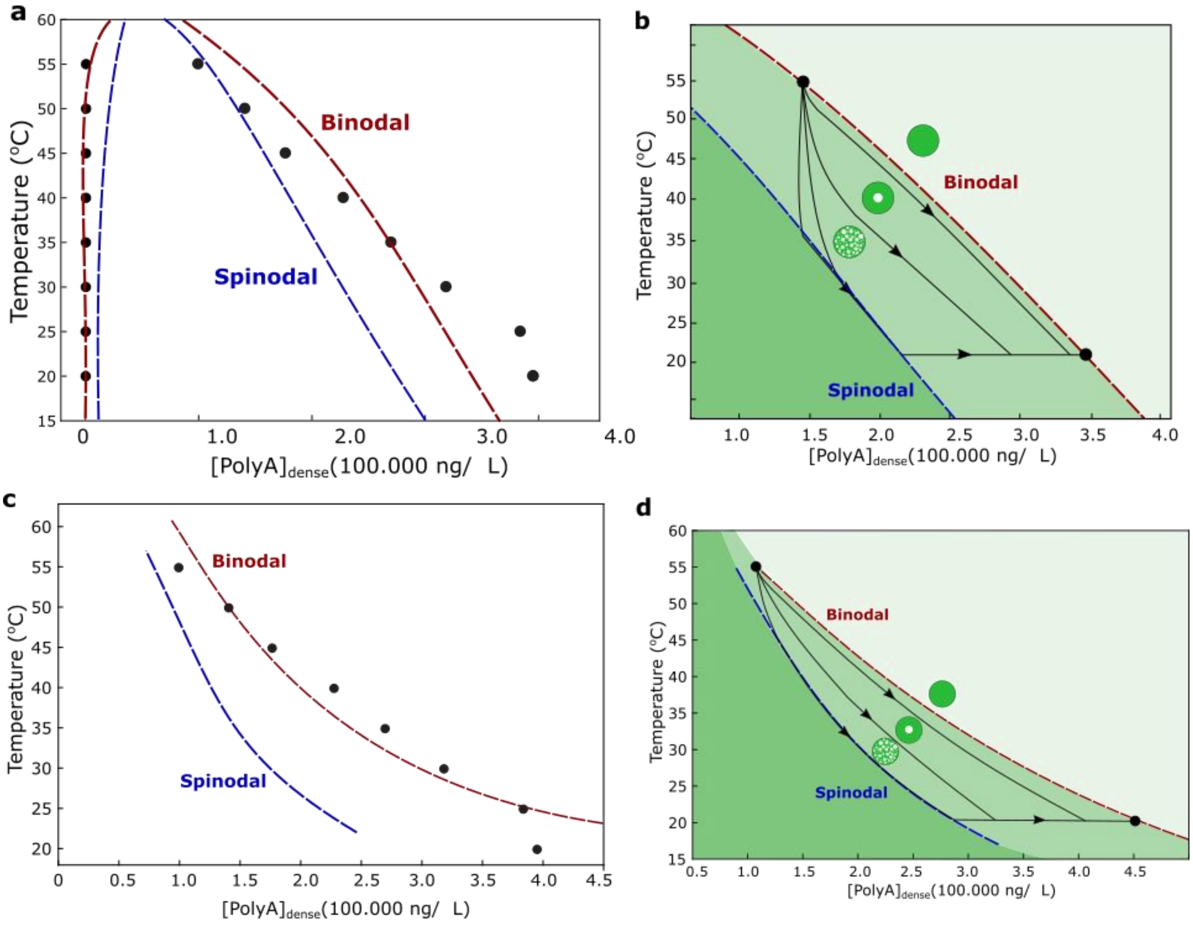


**Supplementary Fig. 7: Quantified binodal and spinodal curve and kinetically arrested phase transitions trajectories.** The dilute and dense phase PolyA concentrations in Supplementary Fig. 1 are fit using a Flory-Huggins model to **a** find the location of the binodal and spinodal curve and **b** consider what trajectories, depending on factors like condensate size, might result in cavity formation (Supplementary text 1). Similarly, the data can also be fit using an electrostatic model to give **c** the location of the binodal and spinodal curve and **d** the associated trajectories. Deviation from the binodal curve can result in the nucleation of dilute phase inside of the condensate.

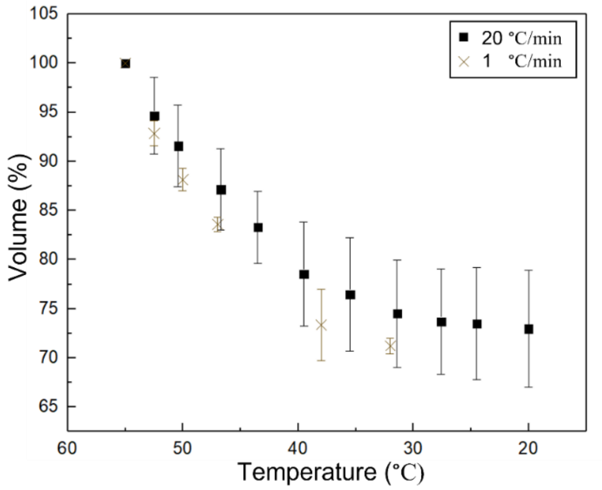


**Supplementary Fig. 8: Volume of PolyA-PEG condensates at different cooling rate.** The volume of condensates cooled at 20 and 1 °C/min was determined by measuring their diameter using confocal microscopy and correcting the temperature using Supplementary Fig. 5. The volume is stated as a percentage of the volume at 55 °C, so that data of 3 condensates, with a slightly different size, could be used to obtain a standard deviation. We observe that there is no significant difference between the sizes of condensates at these rates, indicating that the deviation from the binodal during composition changes is unlikely to be caused by the viscoelasticity of the condensates.

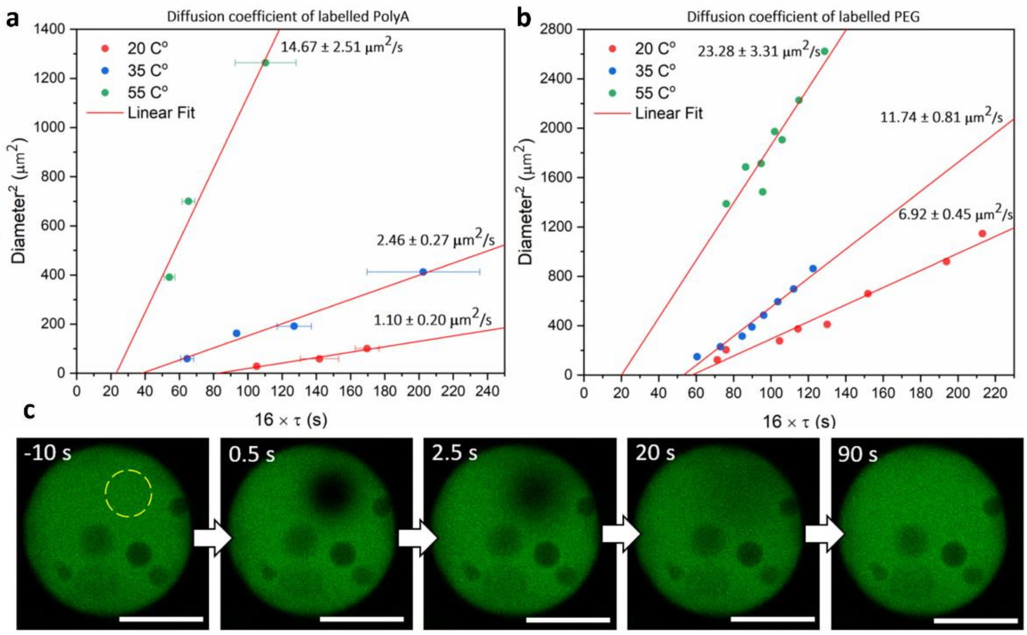


**Supplementary Fig. 9: The diffusion coefficients of PolyA and PEG determined using FRAP.** A circular area in PolyA-PEG condensates with labelled **a** PolyA or **b** PEG are bleached to determine the recovery time of PolyA. The slope of this graph of the diameter squared against 16 times the recovery time is the diffusion coefficient.[23] The diffusion coefficient increases with temperature and the fitted lines for all three temperatures do not pass through the origin, but through $16 \times \tau > 0$. This shows the diffusion was limited by a dense polymer network, rather than purely controlled by Brownian motion[23]. **c** Example of a FRAP experiment in which PEG is bleached (yellow circle) from t = 0 until t = 0.5s. The intensity in the bleached area recovers over time. All scale bars represent 25 μm.

### Supplementary text 1: Binodal and spinodal curve: Flory-Huggins fit

We use two-component Flory-Huggins theory to find the binodal and spinodal curve for our PolyA-PEG system. We approximate our system to be made up of PolyA and solvent. For this system, the free energy density $f$ is [1]

$$f(\phi) = k_B T[\frac{\phi}{N}\ln\phi + (1-\phi)\ln(1-\phi)\,] + \epsilon\phi(1-\phi) \qquad (1)$$

where $k_B T$ is the unit thermal energy, $\phi$ the volume fraction of PolyA, $N$ the effective polymer length and $\epsilon$ the mean-field effective interaction energy. We use a previously established algorithm[1] to determine the location of the binodal. To do this, we use the estimated PolyA concentration (Supplementary Fig. 1) and an

RNA density of 1.6 g/mL[2] to estimate the PolyA volume fraction at the different temperatures. We fit this data to find the parameters $N$ and $\epsilon$ from equation 1 using the algorithm. We fit both parameters $N$ and $\epsilon$, to give $N$ = 1000 and $\epsilon$ = $178k_B \cdot K$ (Supplementary Fig. 7a). By dividing $\epsilon$ by the temperature during our process, which is approximately 300 K, we find that the effective contact energy is roughly $\frac{1}{2}\, k_B T$. The $N$ value is smaller than the number of nucleotides in an PolyA chain, which is between 2100 and 10500. This suggests the effective monomer unit is larger than a single adenosine base, which is sensible considering the chains can have a persistence length longer than the length of single nucleotides. Now that we have estimated the location of the binodal, we use the instability condition $\frac{d^2 f(\phi)}{d\phi^2} = 0$ and solve for $\phi$ to find the spinodal.[3] Supplementary Fig. 7a shows the location of the binodal and spinodal. Supplementary Fig. 7b shows how cavities would be formed during cooling, as the trajectory deviates from the binodal towards the spinodal curve.

**Supplementary text 2: Binodal and spinodal curve: Electrostatic model**

Additionally, we provide fitting of the PolyA concentrations in the dilute and dense phase using an electrostatic model which considers counter ion pressure. This microscopic model describes the competition between the favourable biopolymer interactions and electrostatic repulsion, caused by the entropic cost of placing counterions to screen the charges on the biomolecules in the condensate .[4,5] This compression of screening layers results in the so-called Donnan Pressure,[6] which explains the effects of temperature on the condensate density.

The PolyA-PolyA and PolyA-PEG contact energy are assumed to have a Flory-Huggins form

$$f_{AA} = \varepsilon_{AA} c_A^2 \qquad (3a)$$

$$f_{AP} = \varepsilon_{AP} c_A c_P \qquad (3b)$$

Although in the classical Flory-Huggins picture only cross-interaction terms are present, we can treat the solvent implicitly and write the solvent-solute interaction as solute-solute interaction by invoking the volume constraint $\phi_{solvent} = 1 - \sum_i \phi_{solute}^{(i)}$, where $\phi_{solute}^{(i)}$ and $\phi_{solvent}$ denote the volume fraction of i-th solute and solvent respectively. To further treat the PolyA-PEG interaction, we observe that the PEG concentration in the condensate increases almost linearly with PolyA as temperature is lowered. We thus assume $c_P = a\, c_A + b$ with a and b some constants, and substitute this into equation (1b) to eliminate $c_P$. The term linear in $c_A$ is discarded since it does not affect phase separation calculations. Note this is equivalent to taking a slice of the free energy landscape in the $c_A - c_P$ parameter space to remove one parameter, since we know a priori that the binodal falls onto a straight line in $c_A - c_P$ space and taking this slice will thus not change the binodal we find using the sliced free energy. The result is then simply a free energy quadratic in $c_A$ so we absorb all the interactions and tie-line gradient into a single phenomenological parameter $\varepsilon$, and write $f_{AA} + f_{AP} = \varepsilon c_A^2$.

The electrostatic free energy density is given by

$$f_{ES} = \frac{1}{2}\rho\phi + k_B T\left[c_+ \ln\frac{c_+}{\eta c_s} - c_+ + c_- \ln\frac{c_-}{\eta c_s} - c_- + \eta c_s\right] \qquad (4)$$

Where $\phi$ is the electrostatic potential, $c_\pm$ is the concentration of salt cations/anions, $\rho = e(-c_A + c_+ - c_-)$ is the charge density which has contributions from the phosphate backbones and the mobile salt ions, and $\eta \equiv \frac{V'}{V}$

is the fraction of volume available for salt ions within the condensate. In Eq. 4 the first term represents the Coulomb energy of the system while the second term represents the entropic cost to enrich/deplete mobile cations/anions relative to a reservoir with salt concentration $c_s$. Minimizing Eq. 4 with respect to $c_\pm$ reveals that the salt concentrations follow the Boltzmann distribution $c_\pm = \eta c_s e^{\mp e\phi/k_B T}$ .

Due to the concentrated environment inside the condensate, we employ a "jellium" model in which we approximate the potential as spatially uniform $\phi(x) \simeq \bar{\phi}$ and solve for $\bar{\phi}$ using the condition that the interior of the condensate must satisfy charge neutrality $c_A = c_+ - c_-$.[7,8] Using the Boltzmann relations for the salt concentration this becomes

$$-c_A = 2\eta c_s\, Sinh \frac{e\bar{\phi}}{k_B T} \tag{5}$$

Substituting these into the free energy expression and define the shorthand $x \equiv \frac{c_A}{2(1-rc_A)c_s}$, we get

$$f(c_a) = k_B T c_A \left( \sinh^{-1}(x) - \frac{1}{x}\sqrt{1+x^2} + \frac{1}{2x} \right) + k_B T c_s \tag{6}$$

The last constant term causes the free energy to be 0 for [PolyA] = 0. For $\eta$, we assume a linear relationship with $c_A$ so $\eta = 1 - rc_A$, with $r$ is another parameter to fit.

Combining the derived equations for free energy, the total free energy per volume is given by

$$\frac{F_{tot}}{V} = k_B T \left( c_A \sinh^{-1}(x) - \frac{c_A}{x}\sqrt{1+x^2} + \frac{c_A}{2x} + c_s \right) + \varepsilon c_A^2 \tag{7}$$

To find the binodal in the dense phase for our PolyA model system, we assume that the common tangent construction will give us a y-intercept close to 0, so we will solve $c_A f'(c_A) - f(c_A) = 0$. This gives the relationship between temperature and $c_A$. We solved this for temperature as a function of [PolyA] given:

$$T = -\frac{\varepsilon {c_A}^2}{2k_B c_s\left(\sqrt{1+x^2} - 1\right)} \tag{8}$$

Fitting gives $\varepsilon$ = -1.16 $kJ \cdot L \cdot mol^{-2}$ and $r$ = 0.138 $L \cdot mol^{-1}$. The fitted line resulting from these values and the experimental data points are shown in Supplementary Fig. 7c. Additionally, Supplementary Fig. 7d shows how cavities would be formed if the spinodal is crossed during cooling. The microscopic theory yields binodal and spinodal curves very similar to that of the Flory-Huggins model, and we see again that deviations from the binodal can result in cavity formation.

### Materials and Methods

#### Materials

PolyA (MW 700-3500 kDa), PEG (average MW ~20,000), HEPES, glacial acetic acid, NaCl, KCl, ethanol and mPEG(5k)silane were obtained from Sigma Aldrich. Alexa Fluor™ 647 Carboxylic Acid and RNAase free water was obtained from Thermo Fisher. BactoView™ nucleic acid binding dye (500×) was purchased from Biotium. PEG(20k)-AF647 was purchased from Nanocs. Sylgard 184 Elastomer base and curing agent were bought from Dow Corning Corporation. 18x18 mm glass slides were purchased from Academy. 24x60 mm No.1.5 glass slides were obtained from DWK Life Sciences. Cy3 labelled 100mer ssDNA (CTCACCCACAACCACAAACAATTTAAATAATATTAAATAATATTAATATATTATCGATTAAAT AATAATTAATTAATATTGGTTGGATGGTAGATGGTGA) was purchased from Merck. $(U)_{40}$-Cy3 and $(U)_{40}$-Cy5 for FRET-FLIM were obtained from GenScript. The HzATNP variant 4 antibody was kindly provided by Global Research Technologies Novo Nordisk A/S and the expression is described below. Details on the production of G3BP1 and G3BP1-mEmerald cell lysate are given below as well.

**Fabrication of imaging wells for confocal microscopy**

Holes of 5 mm in diameter were punched in PDMS slabs of ~3 mm height using biopsy puncher. These slabs were plasma bonded to 24×60 mm No.1.5 cover glass slides to create wells. Both the wells and 18×18 mm glass slides used to seal the top of wells were treated with PEG-silane, using a method based on previously reports.[9,10] Briefly, treatment solution was made by mixing 10 mg of PEG(5000)silane with 20 uL of glacial acetic acid and 1 mL of ethanol. Wells and glass slides were treated by placing them in solution for 1 hour at 65 °C and afterwards washing them thoroughly with water. Treated devices are not used after more than 3 weeks since the treatment.

**Making condensates samples**

PolyA-PEG condensates are prepared by mixing stock solution such that we obtain 2000 ng/uL PolyA, 2.5× BactoView™ nucleic acid binding dye, 5 w/w% 20.000 PEG (optionally, of which 0.1% is 20.000 PEG-AF647), 750 mM KCl, 50 mM HEPES at pH = 7.3 in RNAase free water. The PolyA concentration is determined with a nanodrop machine by measuring the absorbance at 260 nm before the addition of the dye.

Reconstituted stress granules were prepared as described previously[11]. Briefly, stress granules were formed by mixing cell lysate containing G3BP1-EGFP with recombinant G3BP1, resulting in a solution with 50 µM recombinant G3BP1 and 1.38 mg/mL protein concentration from the cell lysate. In our experiments, PolyA and 2.5 x BactoView™ dye is added. Notably, BactoView™ stains all nucleic acids, not just the PolyA.

Antibody DNA condensates were made by mixing 5% labelled AF647 HzATNP variant 4 antibody with 100mer Cy labelled ssDNA and buffer to obtain: 6 µM antibody, 18 µM DNA with 15 mM NaCl in 2 mM HEPES buffer at pH = 7.4.

**Confocal imaging**

A Leica Stellaris 5 confocal microscope (confocal fluorescence imaging) (white light laser) microscope equipped with a 10× Nikon 0.3 NA or a 63× oil immersion Leica 1.4 NA is used for imaging. A TS102SI Instec rapid heating and cooling stage is used to control the temperature and change it at desired rates. Qualitative confocal images like the ones in figures 1 and 5 are taken in analog mode. The intensity is then compared to a reference sample, containing PolyA, dye and PEG, to control for the influence of temperature on the brightness of the sample. A similar reference sample is used for the quantitative studies (Fig. 2), however then the photon counting mode was used. To determine the temperature of the sample in comparison to the temperature of the temperature control stage (Supplementary Fig. 5), we measured the intensity of a solution of AF647 (10 uM) in the presence of PolyA-PEG condensates. The intensity of the AF647 scaled linearly with temperature and was thus used to determine the temperature of samples during cooling. For FRAP (Fig. 3), a 488 agon laser at 100% power is used to bleach disk of different area sizes. The recovery of fluorescence over time in the FRAP kymograph is fitted with a single exponential to obtain the half life time τ using the built-in software. FRET-FLIM was performed using the TauSense mode. At each condition, the determined lifetime was average of the lifetime in condensates of at least 10 images. The standard deviation of the lifetimes for the pixels within an image was of less than 0.03 ns. Both the lifetime of just the donor and the donor in presence of the acceptor are measured at each temperature to determine the FRET efficiency E.

Only the images in Supplementary Fig. 10 were taken on an LSM 880 (Zeiss) microscope by exciting using an Argon multi-line 35 mW 488 nm laser (3 mW max at focal plane). The resulting fluorescence was collected using a 63× Plan-Apochromat 1.4 NA oil objective (Zeiss) and detected on a 34-channel spectral array detector in the 480 nm – 580 nm range. Samples were imaged with a Definite Focus module (Zeiss) employed for thermal drift correction and ZEN Black v2.3 (Zeiss) software used for acquisition.

Further analysis of pictures was performed using Fiji.

**Production of HzATNP variant 4**

HzATNP variant 4 antibody was expressed according to previously reported protocols.[12,13] After expression, all antibody samples were stored in aliquots at roughly 5-10 mg/mL in 2mM HEPES Buffer pH = 7.4 with 15mM NaCl. For the labelling the antibody (30µL, 1.1 nmol, 1 equiv.) was diluted further with NaHCO3 (65µL) followed by addition of Alexa Fluor 647 (in DMSO, 1 equiv.) to give a total volume of 96.6 µL, followed by incubation for approx. 1 h at RT. under protection from light. Purification was performed by buffer exchanging to 2mM HEPES Buffer pH = 7.4, 15mM NaCl with 10kDa molecular weight cut-off Amicon Ultra-0.5 centrifugal filter units.

**Production of G3BP1**

Plasmid containing His-Sumo tagged G3BP1-FL was transformed into competent *E. coli* BL21(DE3) (NEB). A single colony was used to inoculate 5.0 mL of LB media containing kanamycin and incubated overnight at 37 °C. The starter culture was subsequently used to inoculate 10 litres of LB media containing kanamycin and incubated at 37 °C until the Absorbance at 600 nm reached 0.6 at which point the G3BP1 expression was induced by 0.6mM IPTG and the cultures were incubated at 16 °C overnight. Cells were harvested by centrifugation at 5000 rpm for 20 minutes. Cell pellet was resuspended in Buffer A (50 mM Tris, 200 mM NaCl, 1 mM DTT pH=8.0) plus protease inhibitor cocktail and subjected to high pressure cell lysis using Constant Pressure Cell Disruption System (Constant systems. Ultracentrifugation at 100.000 × g was employed to clarify the cell lysate prior to loading onto a 20 mL Ni-Sepharose Advance resin containing gravity column (Bioserve). His-SUMO tagged G3BP1 was purified using standard Ni-affinity protein purification protocol which included a wash step in Buffer A containing 25 mM Imidazole and an elution step in Buffer A containing 500 mM Imidazole. The column eluates were run on an SDS-PAGE and the fractions containing the protein were pooled, mixed with His-tagged ULP protease for the cleavage of the His-SUMO tag and dialysed in buffer A overnight at 4 °C. Cleaved protein was further treated with 0.1 mg/mL RNase and DNase by incubating it at 37 °C for 15 minutes. Post incubation, protein sample was diluted using Buffer A without NaCl to lower the salt concentration to 50 mM before loading it on a SP-Sepharose (Cytiva) ion-exchange column. Protein was further purified by running a salt gradient and the fractions containing the protein were pooled, concentrated and subjected to the gel filtration step using a Superdex-200 Increase column (Cytiva) in the storage buffer (50 mM HEPES, 400 mM NaCl, 1 mM DTT, pH = 7.5). Protein fractions containing G3BP1 were assessed by SDS-PAGE and the pure fractions were pooled and concentrated to 15 mg/mL. Protein was aliquoted and snap frozen in Liquid Nitrogen for all subsequent assays.

**Cell Line Generation for G3BP1-mEmerald cell lysate**

To generate the G3BP1-mEmerald HeLa line, a PiggyBac transposon-transposase system was used. Initially, a vector was designed using Gibson assembly of the G3BP1 transgene into a custom mEmerald-PiggyBac vector (available upon request). Cotransfection of the G3BP1-mEmerald-PiggyBac vector and a transposase (TransposagenBio) plasmid into HeLa cells (ATCC) was followed by blasticidin selection to isolate a polyclonal G3BP1-mEmerald population. Transfections were performed using FuGene (Roche) at a 1 µg DNA: 3 µl FuGene ratio. Cells were maintained at 37 °C and 5% $CO_2$ in Dulbecco's Modified Eagle Medium (DMEM, Gibco) supplemented with 10% fetal bovine serum (FBS, Gibco).

**Generation of G3BP1-mEmerald cell lysate**

The generation of cell lysates from G3BP1-mEmerald HeLa cells and the subsequent generation of lysate granules have been described previously[11]. Briefly, cells were grown to 100% confluency in a 10cm dish and harvested in 5 mL PBS. All steps were performed at room temperature. Collected cells were spun at 500 × g for 5 minutes with the supernatant aspirated and discarded. Cell pellets were stored at -80 °C until ready to use. To prepare cell lysates, pellets were thawed for 2 minutes and resuspended in 250 µL lysis buffer containing 50mM TRIS pH7.0 (Sigma), 0.5% NP40 (Thermo), 0.025× mini protease inhibitor tablet (Roche) and 40× murine RNase inhibitor (NEB). After a 3-minute incubation period lysates were spun at 24.000 × g for 5 minutes to remove nuclei and cell debris. The supernatant was retained to produce reconstituted stress granules when mixed with unlabelled G3BP1.
